# Top-down predictions of visual features dynamically reverse their bottom-up processing in the occipito-ventral pathway to facilitate stimulus disambiguation and behavior

**DOI:** 10.1101/2021.10.12.464078

**Authors:** Yuening Yan, Jiayu Zhan, Robin A.A. Ince, Philippe G. Schyns

## Abstract

The prevalent conception of vision-for-categorization suggests an interplay of two dynamic flows of information within the occipito-ventral pathway. The bottom-up flow progressively reduces the high-dimensional input into a lower-dimensional representation that is compared with memory to produce categorization behavior. The top-down flow predicts category information (i.e. features) from memory that propagates down the same hierarchy to facilitate input processing and behavior. However, the neural mechanisms that support such dynamic feature propagation up and down the visual hierarchy and how they facilitate behavior remain unclear. Here, we studied them using a prediction experiment that cued participants (N = 11) to the spatial location (left vs. right) and spatial frequency (SF, Low, LSF, vs. High, HSF) contents of an upcoming Gabor patch. Using concurrent MEG recordings of each participant’s neural activity, we compared the top-down flow of representation of the predicted Gabor contents (i.e. left vs. right; LSF vs. HSF) to their bottom-up flow. We show (1) that top-down prediction improves speed of categorization in all participants, (2) the top-down flow of prediction reverses the bottom-up representation of the Gabor stimuli, going from deep right fusiform gyrus sources down to occipital cortex sources contra-lateral to the expected Gabor location and (3) that predicted Gabors are better represented when the stimulus is eventually shown, leading to faster categorizations. Our results therefore trace the dynamic top-down flow of a predicted visual content that chronologically and hierarchically reversed bottom-up processing, further facilitates visual representations in early visual cortex and subsequent categorization behavior.

---

Since Helmholtz’ “unconscious inferences,” vision scientists have worked with the hypothesis that what we visually perceive is in part influenced by the bottom-up sensory input, but also by top-down expectations of what this input might be [1, 2]. Expectations predict the upcoming visual information [3-5], thereby facilitating its disambiguation from the noisy input [6, 7] to then speed up behavior [8]. Advances in neuroimaging revealed that coordinated networks of brain regions process visual information bottom-up (feedforward) and/or top-down (predictive, feedback) [1, 9]. In bottom-up mode, left and right occipital cortices represent respectively the right and left visual hemifield impinging the retina [10]. These contra-lateral representations then transfer (including across hemispheres) up to the ventral and dorsal regions where they are merged to implement “vision for categorization” and “vision for action”, respectively [12-15]. A convergence of formal models [3, 4, 16] and empirical evidence [17] suggest that the top-down mode should reverse the bottom-up mode. Predictions originating from memory would travel from the top of the “vision for categorization” pathway deep in the right fusiform gyrus, down to the left vs. right occipital cortices, depending on the visual hemifield where the visual features are expected. However, the specific dynamic neural mechanisms and brain regions that propagate such visual predictions down the visual hierarchy to facilitate behavior remain unclear. Here, to clarify the relationships between memory prediction, behavior and the top-down and bottom-up processing of visual information, we addressed three key questions:

1. Does the top-down prediction of a stimulus feature improve its subsequent behavioral categorization?
2. Does the top-down predictive process reverse the bottom-up feedforward processing of a stimulus feature in the visual hierarchy?
3. Does this reversal in turn facilitate the feedforward processing of the upcoming stimulus, when shown, including its behavioral categorization?

## Results

To control predictions, we used the three-stage cueing design depicted in Figure 1A and 1B (see also *Methods, Cueing-Categorization*). On each trial, a spatial cue at Stage 1 (a green dot briefly displayed left vs. right of a central fixation cross, cf. Posner cueing [18]) predicted the visual hemifield location (left vs. right) of an upcoming Gabor patch (henceforth, Gabor) with 100% validity, followed by a 1-1.5s blank screen. At Stage 2, on informative trials an auditory cue (a 250ms sweeping tone at 196 Hz vs. 2217 Hz) predicted the Spatial Frequency content (SF, Low vs. High) of the upcoming Gabor stimulus with a 90% validity (henceforth, refer as the “predicted” trials), followed by another 1-1.5s blank interval. On uninformative, “non-predicted” trials (33% of Stage 2 trials), a 622 Hz neutral auditory cue had no association with LSF or HSF. Finally, at Stage 3, the actual Gabor stimulus appeared in the participant’s left vs. right visual hemifield for 100ms. Each participant (N = 11, see *Methods, Participants*) categorized the Gabor SF as quickly and accurately as they possibly could without feedback (i.e. 3-AFC, with responses “LSF” vs. “HSF” vs. “don’t know”). We concurrently recorded the participant’s dynamic brain activity with MEG and reconstructed it on 12,773 sources, see *Methods, MEG Data Acquisition*.

**Figure 1.**
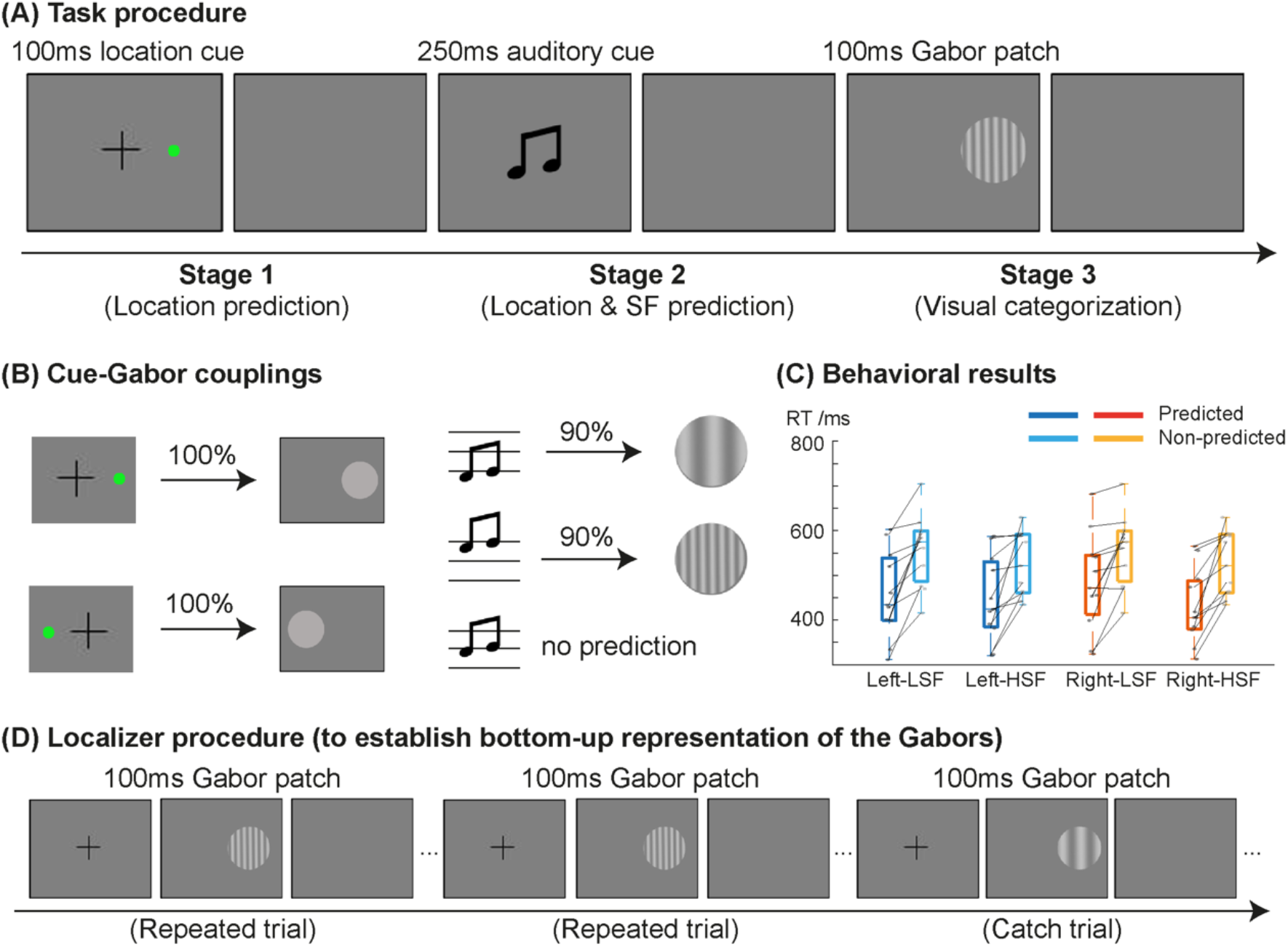
Experimental design and behavioral results. **(A) Task procedure**. Each trial started with a 500ms fixation, followed by Stage 1 where a 100ms green dot (i.e. location cue) that predicted the location of the upcoming Gabor patch, followed by 1000-1500ms blank screen with jitter. At Stage 2, a 250ms sweeping sound (i.e., SF cue) predicted the LSF vs. HSF content of the upcoming Gabor, followed by a 1-1.5s blank screen with jitter. At Stage 3, the Gabor was presented for 100ms. Participants categorized the SF of the Gabor, followed by a 750-1250ms inter-trial interval (ITI) with jitter. **(B) Cue-Gabor couplings**. At Stage 1, the left vs. right location cue predicted the left vs. right location of the upcoming Gabor with 100% validity. At Stage 2, the 196 Hz vs. 2217 Hz informative auditory cue predicted the Gabor LSF vs. HSF contents with a validity of 90%; a 622 Hz neutral auditory cue served as non-prediction control on 33% of the trials, (i.e. 50% probability of LSF or HSF). **(C) Behavioral results**. Boxplots show that the prediction (i.e., informative cueing, dark blue/orange) sped up median LSF vs. HSF Gabor categorization RTs in each presentation condition, compared with non-prediction (i.e., neutral cueing, light blue/orange). Black dots (vs. light grey dots) indicate the per-participant median categorization RTs in predicted (vs. non-predicted) trials, linked to indicate non-predicted RT increases in each individual participant replication. **(D) Localizer procedure (to establish bottom-up representation of the Gabors)**. Prior to the cueing experiment, we ran each participant in an independent MEG localizer. Each trial started with a 500ms central fixation, followed by a 100 ms Gabor, then a 750ms ITI blank screen. In each block of 20 trials (i.e. one of left-LSF, right-LSF, left-HSF, right-HSF), at the fixed location (left or right presented), participants passively viewed 20 Gabors at the same SF. 1 catch trial presented one different-SF Gabor that we instructed participants to detect in each block.

We now address the three questions discussed earlier about the process that predicts specific stimulus feature information (left or right visual field x LSF or HSF Gabor) for categorization behavior.

### 1. Do Gabor predictions improve Gabor categorization behavior?

Valid predictions (i.e., the 90% valid trials with informative cueing) did indeed improve categorization accuracy compared to non-predicted (neutral cue) trials, on average by 2.58% (96.9% vs. 94.3%), *F(1,86)*=22.5, *p*=0.0008, and sped up Reaction Times (RTs), on average by 87.7ms (454.4ms vs. 542.1ms), *F(1, 86)*=20.8, *p*=0.001. RT improvements applied to each condition of presentation (all condition significant, see Figure 1C and Supplemental Table S1 and *Methods, Cueing improves Behavior*) and to each individual participant, obtaining Bayesian population prevalence [19] of 11/11 = 1 [0.77 1] (Maximum A Posteriori (MAP) [95%, Highest Posterior Density Interval (HPDI)], see Supplemental Table S2).

### 2. Does the top-down predictive process reverse the bottom-up hierarchical processing of a Gabor?

There is evidence that the temporal order of processing could reverse between perception and memory predictions of full pictures [17]. However, it remains unclear specifically where (in which brain regions), when (in which time windows) and how specific visual features are predicted. Here, we controlled stimulus feature contents (i.e. left vs. right presentation location; LSF vs. HSF) to explicitly test whether the representation of the same stimulus features reversed between their top-down predictions and their bottom-up processing (schematized in Figure 2A). We did so using a decoder (a cross-validated supervised classification algorithm) to compare the top-down prediction of each Gabor (at Stage 2 of the cueing experiment, i.e. auditory cueing of LSF vs HSF, cf. Figure 1A) to its feedforward bottom-up representation (from a localizer run for each participant prior to the cueing experiment, cf. Figure 1D).

**Figure 2.**
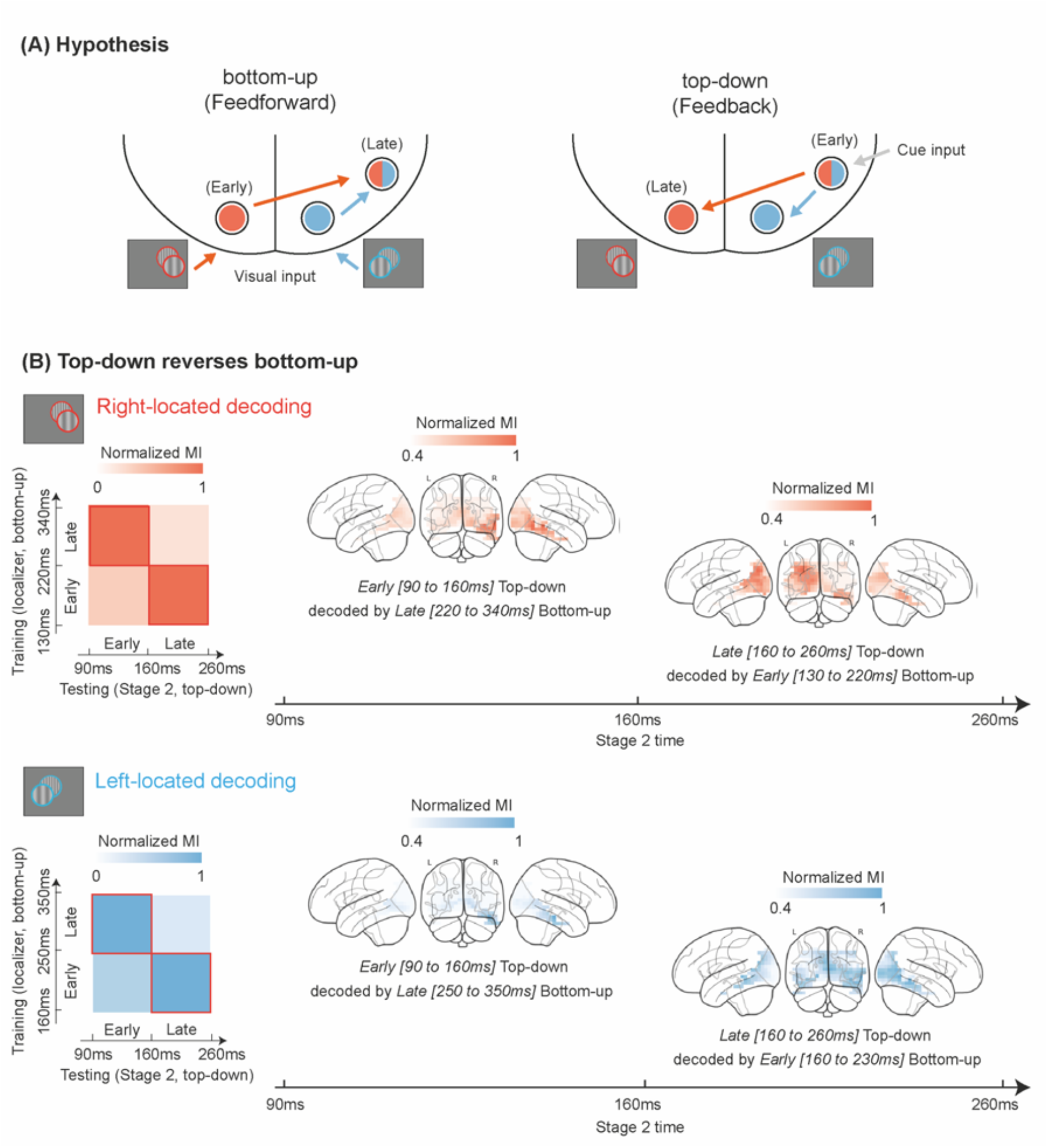
Top-down predictions of left vs. right presented, LSF vs. HSF Gabors reverse their bottom-up processing in the visual hierarchy. **(A) Hypothesis**. We hypothesized that top-down/feedback predictions of Gabor LSF vs. HSF contents dynamically reverse their bottom-up/feedforward processing in the visual hierarchy. That is, the Early Top-Down neural responses at Stage 2 are decoded by the Late Bottom-Up decoder (learned from the MEG localizer), whereas Late Top-Down neural responses at Stage 2 are decoded by the Early Bottom-Up decoder. This would indicate that top-down predictions reactivate LSF vs. HSF Gabor representations, initially in the right fusiform gyrus, where their bottom-up representation ends, to then move to the occipital cortex contra-lateral to expected presentation, where their bottom-up representation starts [14]. **(B) Top-down predictions reverse bottom-up representations in the visual hierarchy**. For right (color-coded in orange) and left (color-coded in blue) located Gabor trials, the 2 × 2 matrix reveals the predicted (cf. A) group averaged cross-decoding performance between bottom-up representation of Gabors (X axis, as training set) in the MEG localizer and their top-down predictions (Y-axis, as testing set) at Stage 2 of the prediction experiment. We quantified decoding performance as MI(<decoder value (continuous values as criteria to classify decoded LSF vs. HSF); ground truth LSF vs. HSF>) and down-sampled time to a 2 × 2 matrix that averaged MI along the Y-axis (separately for right/left-located Gabor trials, with bottom-up intervals assigned to [∼130ms/160ms – 220ms/250ms] and [∼220ms/250ms – 340ms/350ms] time windows) and X-axis (with top-down intervals assigned to [∼90ms – 160ms] and [∼160ms – 260ms] time window). The 2 × 2 matrices represent the group-level mean performance of significant decoding (FWER, *p*<0.05, one-tailed), normalized within each decoder (i.e., normalized across each row). This highlights the reversal whereby the late bottom-up classifier better decodes the early top-down responses, and the early bottom-up classifier better decodes the late top-down responses. <u>Brain plots</u> show the sources that contribute to the classification during the top-down stage. To reveal them, we computed MI(<decoder values; Stage 2 MEG>) along the occipito-ventral pathway (i.e. in lingual gyrus, cuneus, inferior occipital gyrus, middle occipital gyrus, superior occipital gyrus and fusiform gyrus). Source color is the dot product of the time × time decoding performance and the time × time MI(<decoder values; Stage 2 MEG). The brain sources show that top-down LSF vs. HSF predictions reverse their bottom-up representations in the visual hierarchy, starting in the right fusiform gyrus [∼90 – 160ms] post auditory cue at Stage 2 for both right (orange) and left (blue) predicted Gabors and ending in the contra-lateral occipital cortex [∼160 – 260ms].

Specifically, we first trained decoders to classify the Gabor LSF vs. HSF from the MEG localizer responses, separately for left and right Gabor presentations and every 2 ms between 0 and 500 ms post onset (within-participant Multivariate Pattern Analysis [20], cross-validated, FWER [21], *p<*0.05, one-tailed, see *Methods, Multivariate Decoding Analysis, Bottom-up Cross-Validation*). Then, we applied these bottom-up decoders to classify the LSF vs. HSF contents of the top-down predicted Gabor at Stage 2, when the integrated spatial and informative auditory cues enable full prediction of the upcoming stimulus (i.e. its LSF vs. HSF, separately for left vs. right location, cf. Figure 1, *Methods, Multivariate Decoding Analysis, Top-down Cross-decoding*).

We reasoned that if top-down reversed bottom-up Gabor representations, a common pattern of neural activity would instantiate the LSF vs. HSF contents in both the bottom-up decoders prior to the experiment and in the top-down predictions at Stage 2. Note that this common pattern of neural activity cannot result from the evoked responses to the auditory cues, because these were only presented in the prediction experiment at Stage 2 and not in the localizer task used to train the decoder weights. That is, the decoder’s weights could only learn from the visual LSF vs. HSF Gabor presentation, separately for left vs. right presentations.

Figure 2B reports such common patterns between bottom-up and top-down Gabor representations, but in a temporally reversed order. Group averaged top-down predictions of LSF vs. HSF Gabor contents at Stage 2 reverse their bottom-up representations in the hierarchical occipito-ventral pathway (FWER [21], *p<*0.05, one-tailed). That is, reversal starts in the deep sources of the right fusiform gyrus, near auditory cortex area A1, where the auditory cue is processed at (∼90-160 ms post auditory cue, see Figure 1), for both left and right predicted Gabors. Importantly, following this, SF predictions then diverge in the occipital sources contra-lateral to the predicted location of the upcoming Gabor (∼160-260ms, Figure 2B, brain plots). Contra-lateral occipital sources representations of late predictions were significantly different between left and right hemispheres as a function of right and left predicted locations—i.e. group-level ANOVA, 2 (left vs. right prediction) by 2 (right vs. left occipital cortex), *F*(1, 34) = 5.65, *p* = 0.02), an effect replicated in 8/11 participants (Bayesian population prevalence = 0.73 [0.42 0.91] (MAP [95% HPDI], see Supplemental Table S4 and Supplemental Figure S2).

In sum, our analyses showed that the top-down predictive process reverses the bottom-up processing of specific stimulus features in the visual hierarchy. Specifically, predictive cues for left vs. right upcoming Gabors at Stage 2 reactivate the LSF vs. HSF contents represented in right fusiform gyrus sources. These contents then flow down the visual hierarchy to the left vs. right occipital cortex location contra-laterally predicted by the visual cue at Stage 1.

### 3. Do top-down predictions facilitate the subsequent processing of Gabor stimuli for categorization behavior?

Having shown that specific LSF vs. HSF Gabor contents are predicted top-down and contra-laterally at Stage 2, we now show that these predictions serve to improve the initial representations and later categorizations of the Gabors when the stimulus is finally shown at Stage 3 (cf. Figure 1). This is important to understand the specific effects of valid predictions (i.e., valid informative cueing, as opposed to non-predictive) on the initial neural representations of the same physical Gabor stimulus when it is shown, and how these in turn facilitate behavior at a later stage. To preview the results, we will show:

1. That LSF vs. HSF predictions at Stage 2 improve early LSF vs. HSF representations of the Gabor shown at Stage 3 (see schematic brain under Stage 3 in Figure 3).
2. That valid predictions at Stage 2 also modulate premotor cortex activity at Stage 3, speeding up RTs on predicted trials (see schematic brain under RT in Figure 3).

**Figure 3.**
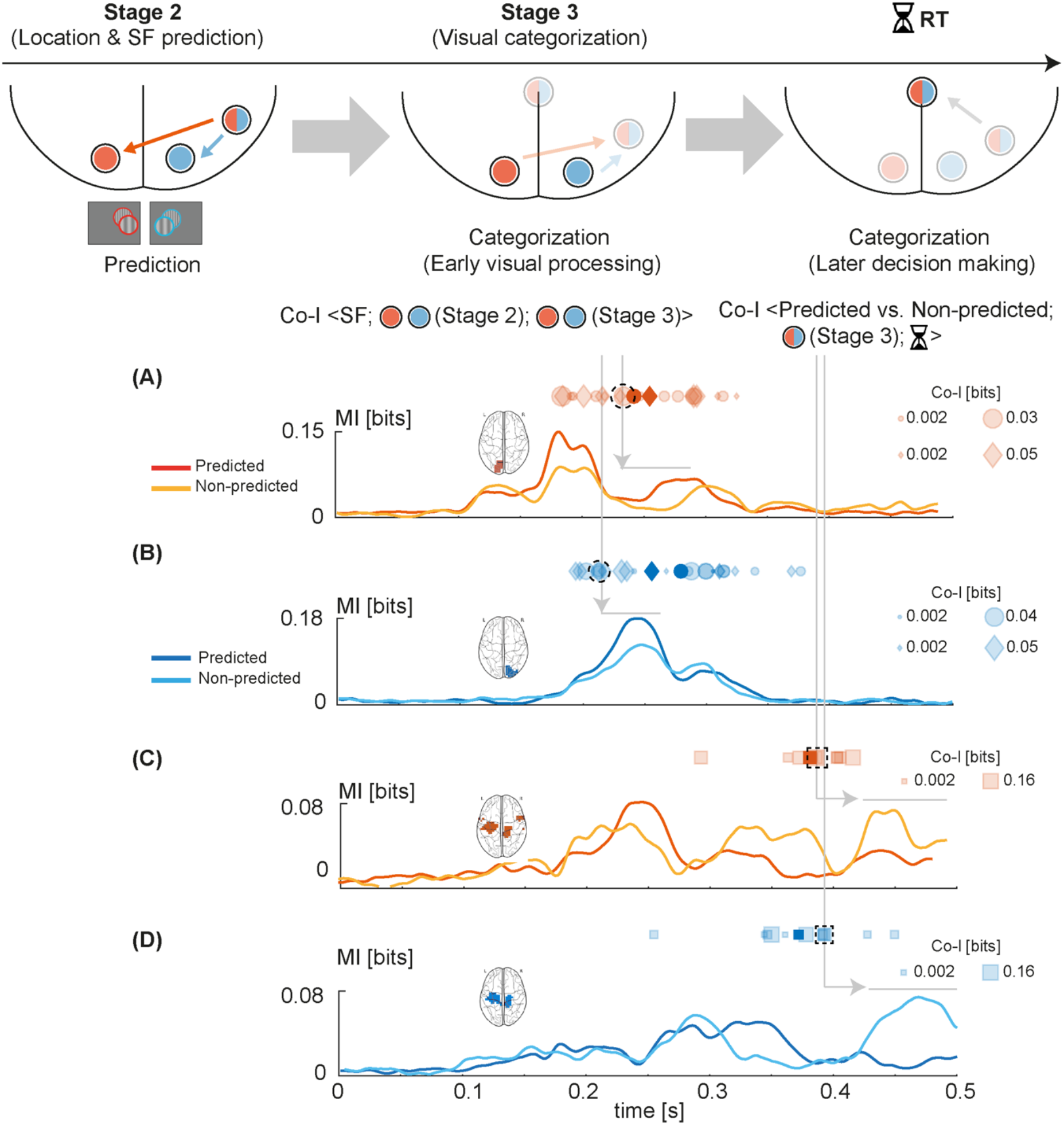
Gabor predictions facilitate feedforward processing of the upcoming Gabor and behavioral RTs. <u>Schematic panels</u> illustrate the logic of the analyses of the effects of Stage 2 prediction representation on Stage 3 Gabor and categorization RTs. Specifically, we tested at Stage 3 whether the contra-lateral occipital cortex representation of the Gabor stimulus SF overlaps with its Stage 2 prediction—i.e. computed as the representational overlap Co-I (LSF vs. HSF; Stage 2 MEG; Stage 3 MEG). Panels A and B underneath present these results Relatedly, we tested whether predicted vs. non-predicted SFs at Stage 2 modulate both Stage 3 premotor cortex activity and RT— i.e. computed as Co-I (Predicted vs. Non-predicted; Stage 3 MEG; RT). Panels C and D present these results. <u>Panels A and B</u> For left-presented (color-coded in blue) and right-presented (color-coded in red) Gabors, we separately averaged left and right occipital sources at the top 25% of the Co-I(LSF vs. HSF; Stage 2 MEG; Stage 3 MEG) distribution (A, B). The horizontal scatters show these Co-I values at their peak time, color-coded as opaque for the group average and as transparent for each individual participant. Circles (vs. diamonds) represent the Co-I peak time between Stage 3 occipital sources representation of LSF vs. HSF with Stage 2 contra-lateral occipital cortex (vs. Stage 2 rFG). The MI curves show the maximum MI (LSF vs. HSF; Stage 3 MEG) for a typical participant. The participant’s dashed Co-I value (in the horizontal scatter) is linked with grey arrows to the MI curves of predicted vs. non-predicted trials. The MI curves show that Gabor SF representation is higher in predicted than non-predicted trials (FWER, *p*<0.05, two-tailed, significant time windows are indicated by solid grey lines) in contra-lateral occipital cortex at Stage 3, following the peak of representation overlap with Stage 2 SF prediction. <u>Panels C and D</u> We similarly averaged the premotor cortex sources at the top 25% of the Co-I (predicted vs. non-predicted; Stage 3 MEG; RT) distribution. The horizontal scatters here show these Co-I values at their peak time, color-coded as opaque squares for the group average and as transparent squares for each individual participant. The later MI curves show that SF representation is reduced in premotor cortex for predicted trials, following the peak of co-representation of predicted vs. non-predicted trials in Stage 3 MEG and RTs (FWER, *p*<0.05, two-tailed, significant time windows indicated by solid grey lines).

#### LSF vs. HSF predictions improve early occipital Gabor representations at Stage 3

Here, we show that valid Stage 2 cueing LSF vs. HSF improves LSF vs. HSF representation of the Gabor stimulus when it is shown at Stage 3. To do so, we proceed in two steps. First, we show that the Gabor SF feature is similarly represented in neural activity (i.e. MEG sources) between its prediction at Stage 2 and when it is physically shown at Stage 3. With predicted trials, we computed this per-trial representational overlap with the information theoretic Co-Information(<LSF vs. HSF; Stage 2 MEG sources; Stage 3 MEG sources>) [22], using rFG and contra-lateral OCC sources at Stage 2 (cf. Figure 2B) and the sources of 6 main ROIs of the occipito-ventral pathway primarily involved with vision-for-categorization [11] (spanning contra-lateral occipital cortex, right fusiform gyrus, temporal lobe, parietal lobe, frontal lobe and pre-motor cortex) at Stage 3 (see Supplemental Figure S3), see *Methods, Representation Overlap*. We found that Gabor SF representations overlapped mostly on contra-lateral OCC sources between Stages 2 (when LSF vs. HSF is predicted) and 3 (when LSF vs. HSF is shown). The round and diamond scatters above Figure 3A and B, indicate the peak amplitudes (highest 25%) and peak times of this representational overlap of LSF vs. HSF Gabor representations into MEG source activity.

Having demonstrated that predictions of Gabor LSF vs. HSF overlap with their actual representations when shown, we turn to the second step of our demonstration. Here, we show that with this overlap, the Gabor LSF vs. HSF prediction at Stage 2 improves the representation of the LSF vs. HSF of the actual stimulus at Stage 3 (compared with non-predicted trials). To do so, we compare the representation of LSF vs. HSF on contra-lateral occipital sources with the peak overlap, for predicted (i.e., valid informative cueing) vs. non-predicted (i.e., neutral cueing) trials, and separately for left vs. right Gabor presentation—i.e. computed as MI(<LSF vs. HSF; Stage 3 MEG>) for predicted vs. non-predicted trials, and for left vs. right Gabor presentations [22]. Figure 3 presents the results of these comparisons. Plots A (for right presented Gabor) and B (for left presented Gabor) show stronger SF representation for predicted trials (darker hues) on the sources that maximally overlapped with prediction (i.e. top 25% of the Co-Information above) in left (vs. right) color-coded OCC, just after the representational overlap peak (data from a typical participant; individual overlap scatter highlighted with the dash frame and grey arrow). Thus, valid prediction at Stage 2 did indeed significantly enhance Stage 3 representation of the LSF vs. HSF Gabors in contra-lateral OCC (FWER, *p*<0.05, two-tailed), within a 50ms time window following the overlap peak time. Supplemental Figure S4 presents the results for each individual participant, replicating the effect for left located trials in 8/11 (Bayesian population prevalence = 0.73 [0.42 0.91] (MAP [95% HPDI]) and 9/11 in right located trials (Bayesian population prevalence = 0.81 [0.53 0.96] (MAP [95% HPDI]).

In sum, we have shown that Stage 2 SF predictions of the upcoming left vs. right Gabor enhance the early representation of this stimulus feature specifically in contra-lateral right vs. left occipital sources at Stage 3, when the Gabor stimulus is shown.

#### Valid predictions modulate pre-motor cortex activity and speed up categorization

Here, we further show that Stage 2 predictions also modulate Stage 3 pre-motor cortex activity and subsequent RTs. To do so, we computed the overlap of representation between predicted vs. non-predicted trials at the Stage 3 neural activity of the sources of the 6 ROIs used earlier and behavioral RTs—i.e. as Co-Information (<predicted vs. non-predicted; Stage 3 MEG; RT>), separately for left vs. right-presented trials (FWER-corrected, *p*<0.05, see *Methods, Cue modulates Source Activity and RT*).

In Figure 3, the square scatters above plots C and D indicate the timing and amplitude of highest 25% co-representation of prediction on Stage 3 premotor cortex source activity and RT. This co-representation peaks at ∼400ms post Gabor presentation, speeding up categorization RTs (see Supplemental Figure S3 for whole brain result). To understand this specific effect on pre-motor cortex sources, we compared their LSF vs HSF representation of predicted vs. non-predicted trials— i.e. computed as MI(<LSF vs. HSF; Stage 3 MEG>), for pre-motor cortex sources at the highest 25^th^ percentile of the Co-Information distribution above. Figures 3 C and D present these MI plots, where pre-motor cortex sources maintained their representation of LSF vs. HSF Gabor for a longer duration on non-predicted vs. predicted trials after the co-representation peak (the co-representation scatter of the typical participant is highlighted with the dash frame and grey arrow). This suggests a decision-making latency for non-predicated trials that we replicated in individual participants (see Supplemental Figure S4, replicated for left located trials in 10/11 participants, Bayesian population prevalence = 0.91 [0.64 0.99], MAP [95% HPDI], and for right-located trials in 8/11 participants, Bayesian population prevalence = 0.73 [0.42 0.91], MAP [95% HPDI]).

In sum, we showed in this section that valid predictions shorten the representation of LSF vs. HSF in premotor cortex, incurring faster RTs.

## Discussion

We asked three key questions pertaining to the dynamic mechanisms that predict specific visual contents in the brain (i.e. top-down, feedback processes) and their relations to the representation of these same contents from the stimulus itself (i.e. bottom-up, feedforward processes). Our approach used a three-stage predictive experimental design: At Stage 1, a spatial cue predicted the location (left vs. right) of an upcoming Gabor; At Stage 2, an auditory cue predicted the SF contents of the stimulus; At Stage 3, the Gabor stimulus was presented. With those manipulated predicted contents, we asked three main questions. First, whether top-down predictions of a stimulus from memory improve its subsequent categorization? At behavior-level, we found that valid predictions of Gabors sped up categorization RTs in all participants and improved accuracy in most. Second, we asked whether the top-down predictive processing of a Gabor reverses its bottom-up representation in the visual hierarchy? We found that it did so, with a top-down process located deep in rFG sources to propagate down to the predicted contra-lateral OCC. Finally, we asked whether such predictive reversal facilitates the feedforward processing of Gabor stimulus shown at Stage 3 and speeds up its categorization? Our results showed that predictions overlapped with actual stimulus processing, leading to enhanced early representation of the displayed stimulus in the contra-lateral occipital sources and that predictions shortened pre-motor cortex Gabor representations and in turn sped up RTs. Together, our results support the hypothesis that the dynamic top-down flow of a predicted visual content reverse its bottom-up representation, and facilitates visual representations of the physical stimulus in early visual cortex and subsequent behavior.

### Generalizing from Gabors to other stimulus features in face, object, body and scene categorizations

Our first contribution is methodological, providing the means to isolate the specific dynamic processing of a predicted stimulus feature into the visual hierarchy. Generalizing from our case study with Gabor stimuli to more naturalistic face, object and scene categorization tasks will incur several challenges that we can frame in the context of generally studying the details of information processing [23, 24]. A key challenge is that the features of real-world faces, objects and scenes, may differ across behaviors and participants, rather than being fixed as they were here to Gabor LSF vs. HSF. Such task- or participant-specific features remain to be considered in studies of complex neural representation, processing and categorization, though see [14, 15, 25]. A key constraint of studying information processing is that we need to characterize the information processed as precisely as possible. In complex categorizations, this implies understanding which specific image features are task-relevant and therefore predicted by each participant that performs the same behavioral task, but possibly with different stimulus features [26-29]. Furthermore, we also need to know the behavioral outcomes of the process, that the typical passive viewing, or one-back designs of neuroimaging experiments do not control. For example, a participant will predict a smiling mouth feature to verify that a given incoming face is “happy” but the forehead wrinkles to predict “45 years of age” from the same face. Relatedly, experts will use different features than novices to classify their expert categories and such relative perceptual expertise and associated features generally characterize the relationship between visual cognition and outside world stimuli [30, 31]. A methodological challenge will be to study visual predictions along the hierarchical decomposition of features, from their integrated representation in rFG, to their simplest Gabor representation in V1. This will likely require integration of brain measures that finely tap into the laminar layers of these regions [32], with measures that tap into dynamic flow [33]. In sum, to understand the information processing subtending dynamic predictions in the brain, we need to understand the task behavior that the participant engages with and the stimulus features of the task [34, 35] and then trace predictions of these features and their decompositions across the different hierarchical stages between rFG and V1 [3].

### Generalizing from visual features to features in other sensory modalities

Predictions of stimulus information can in principle occur in any sensory modality. The methodological framework proposed here within the visual modality could straightforwardly extend to other sensory modalities—e.g. auditory—by adapting the design proposed in Figure 1. For example, auditory stimuli can also be decomposed in low-[36] or high-level [37] stimulus properties, for which localizer bottom-up dynamic signatures could be compared with the top-down reactivations of the properties, as we did with our left- and right-presented SF Gabors. From thereon, one could also examine the time-course of predicted multi-modal information in each modality.

To conclude, with a three-stage predictive design, we have traced the dynamic top-down flow of a predicted visual content (left vs. right, LSF vs. HSF Gabor path). We found that bottom-up-reversed predictions improve early visual cortex representation of the shown stimulus and its subsequent categorization. As discussed, our methodology in principle generalizes to predictions in other sensory modalities and from simple to more complex stimulus features and categorizations.

## Methods

### Participants

Eleven participants (18-35 years old, mean=26.8 years old, SD=3.0 years old) participated provided informed consent. All had normal or corrected-to-normal vision and reported no history of any psychological, psychiatric, or neurological condition that might affect visual or auditory function. The University of Glasgow College of Science and Engineering Ethics Committee approved the experiment.

### Stimuli

We used 2 types location cues (green dots) at Stage 1, 3 sweeping sounds as SF cues at Stage 2 and 2×2×3 types (location × SF × orientation) of Gabor patches at Stage 3, as the actual stimuli.

#### Location Cues

We presented the left (vs. right) 1 deg of visual angle diameter green dot at a left vs. right 2 deg of visual angle eccentricity. Participants sat at a 182 cm viewing distance from the screen.

#### SF Cues

Three sweeping sounds started with auditory frequency of 196Hz, 2217Hz or 622Hz, a sweep rate of 0.5 rising octave/second, each for a 250ms total duration.

#### Gabor Stimuli

The left (vs. right)-presented Gabor patch were presented with the diameter of 7.5 visual degree and eccentricity of 12.5 visual degree (left vs. right from the centre), at a 182 cm viewing distance, with LSF (vs. HSF) contents of 0.5 cycle/degree (vs. 1.2 cycle/degree) and one of three randomly chosen orientations (−15 deg, 0 deg, +15 deg). Prior to the task, we calibrated the LSF and HSF Gabor contrast independently for each participant, using an adaptive staircase procedure with target accuracy set at 90%. On each calibration trial, a left (vs. right) green dot presented for 500ms predicted the upcoming left vs. right location of the LSF or (HSF) Gabor patch, itself presented for 100ms. Participant responded “LSF” vs. “HSF” vs. “don’t know” without feedback. We adaptively adjusted LSF vs. HSF as follows:

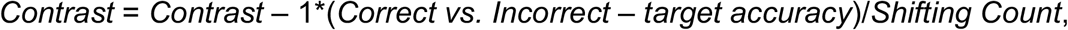

where *Shifting Count* counts the number of direction changes (i.e., increasing to decreasing, or decreasing to increasing). The adaptive staircase stopped when the adjustment step was < 0.01, setting each SF contrast for the Gabor stimuli of this participant in the actual experiment (see Supplemental Table S5 provides each participant’s contrast values).

### Tasks

#### Cueing-Categorization

Each trial started with a central fixation presented for 500ms, followed by three stages (see Figure 1A for task procedure):

At Stage 1, a green dot was presented for 100ms, either left or right of the central fixation, predicting the left vs. right location of the upcoming Gabor with validity of 1. This was followed by a blank screen presented for 1000-1500ms with jitter.

At Stage 2, one of three sweeping sounds was presented for 250ms. The 196Hz (vs. 2217Hz) sound predicted the upcoming LSF (vs. HSF) Gabor (both with .9 validity). The 622Hz sound was a neutral cue that did not have any predictive value. It was followed by LSF vs. HSF Gabors with .5 probability, on 33% of trials).

At Stage 3, the LSF vs. HSF Gabor stimulus appeared at one of the three rotations on the left vs. right screen location for 100ms. The Gabor was either LSF or HSF, with one of three randomly chosen orientations, followed by a 750 to 1,250ms intertrial interval (ITI) with jitter. We instructed participants to respond “LSF” vs. “HSF” vs. “Don’t know” as quickly and as accurately as they possibly could. They did not receive feedback.

The experiment comprised blocks of 54 such trials (see Supplemental Table S6 for the repetitions on each type of stimuli). Participants performed 10-14 blocks in a single day, with short break between blocks. They completed the total of 38-45 blocks over 3-4 days. Participants completed at least 499 trials for each of the 4 (location × SF combination) Gabor patches presentation. Participants learned the correct relationships between the auditory cues and predicted Gabor SF without explicit instructions within two trial blocks. Thus, we removed the first two blocks from all subsequent analyses.

#### Localizer

Prior to the experiment, we also ran an MEG localizer to model the bottom-up processing of each one of 2×2×3 Gabor patches (i.e. location × SF × orientation; Figure 1D illustrates the localizer task procedure). For each Gabor, each localizer trial started with a 500ms central fixation, followed by the Gabor for 100ms, then a blank screen for 750ms ITI. In a block of trials, we repeated the same Gabor for 20 trials in a row. Participants passively viewed the Gabor but had to detect another Gabor (at the same location and orientation) with other SF. Each participant completed 36 such blocks (i.e., 3 blocks per type of Gabor), with block order of “left-LSF-clockwise”, “left-LSF-vertical’, “left-LSF-anticlockwise”, “left-HSF-clockwise”, “left-HSF-vertical’, “left-HSF-anticlockwise”, “right-LSF-clockwise”, “right-LSF-vertical’, “right-LSF-anticlockwise”, “right-HSF-clockwise”, “right-HSF-vertical’, “right-HSF-anticlockwise”, repeated three times.

### MEG Data Acquisition and Pre-processing

We measured participants’ MEG activity with a 248-magnetometer whole-head system (MAGNES 3600WH, 4-D Neuroimaging) at a 508Hz sampling rate. We performed the analysis according to recommended guidelines using the FieldTrip toolbox [38] and in-house MATLAB code.

#### Cueing-Categorization Task

For each participant, we epoched the raw data into trial windows, separately for each stage: Stage 1, -200ms pre-dot onset to 1,000ms post-dot onset (henceforth [-200ms 1,000ms] around onset); Stage 2: [-200ms 1,000ms] around sweeping sound onset; Stage 3: [-200ms 600ms] around Gabor patch onset. Then, we applied 1Hz high-pass filter (5^th^ order two-pass Butterworth IIR filter) to the epoched data, removed the line noise using discrete Fourier transform and de-noised via a PCA projection of the reference channels. We rejected noisy channels with a visual selection and rejected jump and muscle artifacts with automatic detection. We decomposed the output dataset with ICA, identified and removed the independent components corresponding to artifacts (eye movements, heartbeat; 2-4 components per participant).

We then resampled the output data at 512 Hz, low-pass filtered the data at 25Hz (5^th^ order Butterworth IIR filter), specified the time of interest between 0-500ms (post cue at Stage 2; Gabor stimulus at Stage 3), and performed the Linearly Constrained Minimum Variance Beamforming (LCMV) analysis to reconstruct the time series of sources on a 6mm uniform grid warped to standardized MNI coordinate space. Following the above steps, for each participant we obtained single-trial time series of 12,773 MEG sources at a 512Hz sampling rate between 0 and 500ms for Stages 2 and 3. We used these to analyze dynamics information processing in the brain (see Figure 1). Supplemental Table S7 reports the numbers of trials that remained after pre-processing and LCMV analysis.

#### Localizer Task

We applied the same pre-processing pipeline to the MEG localizer, using the epoched data [-200ms 500ms] around Gabor patch onset. The LCMV analysis was then applied 0-500ms post Gabor patch, to reconstruct the source representation of the MEG localizer data. Supplemental Table S7 summarizes the numbers of trials that remained after pre-processing and LCMV analysis.

### Analysis

#### Cueing improves Behavior

At a group-level, we discarded invalid trials and applied a 2 (left vs. right location cues) × 2 (valid informative vs. neutral SF cues) × 2 (LSF vs. HSF Gabor patches) ANOVA on median RTs (excluding incorrect response and outliers) and accuracies (ACC) of valid trials of all participants. We found a significant main effect of valid informative vs. neutral SF cueing on RTs, showing that predicted trials RTs are significantly faster than non-predicted trials (*F(1,86)*=20.8, *p*=0.001); and a significant interaction effect between location cue and Gabor SF (*F(1,86)*=17.4, *p*=0.002). Further analysis showed that this informative vs. neutral cueing effect is significant (*p*<0.05, after Bonferroni correction) for each of the 4 experimental conditions (left vs. right locations × low vs. high SFs), quantified by a paired-sample t-test independently for each condition. For categorization accuracy (ACC), the ANOVA was significant only for valid informative vs. neutral cueing, showing that ACC is significantly higher in in predicted than non-predicted trials (*F(1,86)*=22.5, *p*=0.0008); and a significant interaction between location cue and Gabor SF (*F(1,86)*=13.8, *p*=0.004). Further analysis showed that this effect of SF cue is significant (*p*<0.05, Bonferroni correction) for all but the left-LSF experimental conditions (paired-sample t-test independent for each condition).

We analyzed the RTs (excluding trials with incorrect responses and outlier trials) and ACC of each individual participant. In each participant, we replicated the significant main effect of SF cue on RTs (tested with the 2 × 2 × 2 ANOVA). In each participant, we also examined the effect of SF cue for experimental conditions (independent-sample *t*-tests). Supplemental Table S2 summarizes these ANOVA and independent-sample *t*-tests on RT statistics for each participant. Supplemental Table S3 summarizes the statistics of 2 × 2 × 2 ANOVA and chi-squared test for the accuracy data.

#### Multivariate Decoding Analysis

##### Bottom-up Cross-validation

To model the bottom-up representation of LSF vs. HSF Gabors, we trained decoding classifiers (Linear Discriminant Analysis, LDA) using the MEG localizer, separately for left and right localized Gabors. In each stimulus condition, we segmented the participant’s trials into 5 folds based on stratified sampling and performed a 5-fold cross-validation.

In each iteration, we proceeded in 2 steps:

Step 1: We trained a LDA-classifier ([20], MVPA-Light toolbox in Matlab 2015b) to discriminate LSF vs. HSF Gabors, every 2ms between 0 and 500ms post stimulus, using the MEG sensor data from the trials of 4 folds as training set.

Step 2: We tested the trained LDA classifier at Step 1, every 2ms, with the left-out fold trials, leading to a decision value (i.e. “decoder value”), which was the inner product between the classifier weights and the held-out trial activity at this time point.

Following all 5 iterations, we proceeded to Step 3:

Step 3: To quantify decoding performance on a common scale at each time point, we calculated the Mutual Information (MI, [22]) across trials (concatenating test sets across folds) between this classifier decision value and the true “LSF” vs. “HSF” stimulus label. MI quantifies all the LSF vs. HSF information about the stimulus available from the classifier weights, without forcing a specific discrete prediction [39].

We repeated Step 1-3 10 times and averaged the resulting 10 MI matrices to quantify Gabor decoding over time. To establish statistical significance, we repeated the decoding procedure described 1,000 times with shuffled SF labels, applying threshold-free cluster enhancement (TFCE, *E*=0.5, *H*=0.5, [40]) and using as statistical threshold the 95^th^ percentile of 1,000 maximum values (each taken across the 256 times points per shuffle after TFCE) (i.e., FWER, *p*<0.05, one-tailed, [21]). We repeated the same cross-validation for each individual participant (see Supplemental Figure S1 for significant cross-validation performance for each participant).

##### Top-down Cross-decoding

To determine whether the top-down prediction at Stage 2 comprises SF contents, we used a cross-decoding approach. Specifically, we used the bottom-up SF classifiers trained with localizer data prior to the cueing task, to decode the SF contents at Stage 2, when the stimulus was not shown yet, but we expected the cues to reactivate the stimulus contents. We proceeded in 3 steps, separately for left and right localized stimuli:

Step 1: We trained the bottom-up classifiers every 2ms between 0 and 500ms post Gabor onset, using all MEG localizer trials in each location condition.

Step 2: We tested each bottom-up classifier on its classification of the MEG sensor data at Stage 2, at its specific time point post auditory cue, when prediction should reactivate the cued LSF vs. HSF into MEG source activity. On each trial, the outcome is a 2D (training time × testing time) decision value matrix. To quantify decoding performance, across trials we computed for each combination of training time and testing time the MI between single-trial classification decision value and the true stimulus label (“LSF” vs. “HSF” prediction). A permutation-based threshold (1,000 repetitions), using TFCE to correct for multiple comparison (FWER over 256 × 256 time points, one-tailed, *p* < 0.05) established statistical significance.

Step 3: We down-sampled the decoding performance MI matrix (of dimensions training time × testing time) into a 2×2 matrix that captures the main temporal decoding characteristics. Along the training time axis, we set the early bottom-up time window (i.e., early bottom-up stage) as [147 ± 30ms, 232 ± 42ms] (mean ± standard deviation across participants) and the late bottom-up time window as [232 ± 42ms, 344 ± 48ms], both post localizer Gabor onset. Along testing time axis, we set the early top-down time window (i.e., early top-down stage) as [90ms, 160 ± 20ms] post Stage 2 auditory cue onset, and the late top-down time window as [160 ± 20ms, 260ms]. Note that while these windows were chosen by inspection to visualise the results, our inference corrected FWER over the entire time course at 2ms resolution and is not affected by this choice of bins for visualisation. We averaged the time × time MI matrix (of statistically significant MI values) within the down-sampled time windows, to derive the early-late bottom-up × early-late top-down decoding performance. Finally, within the early and late bottom-up classifier (i.e., Y-axis), we normalized the 2-by-2 matrix across early and late top-down (i.e., X-axis), respectively.

We repeated the above Steps 1 to 3 for each participant to create their down-sampled 2×2 matrix of decoding performance (see Supplemental Figure S2 for each individual result). Figure 2 shows the average of these matrices across participants, separately for the predictions of a left and right located Gabor.

##### Source Representation of Cross-decoding

To visualize top-down reactivation of LSF and HSF Gabor representations at MEG source level, we reconstructed the occipito-temporal sources that cross-decode the Gabor predictions in 3 steps, separating left and right located trials.

Step 1: Using the Stage 2 single-trial decision matrix above (see *Top-Down Cross Validation Step 2*), we computed MI in each cell between the single-trial decision value and Stage 2 single-trial source activity at this specific time point. We confined this analysis to the occipito-temporal pathway involved with vision-for-categorization [14], with regions of interest (ROIs) lingual gyrus, cuneus, inferior occipital gyrus, middle occipital gyrus and superior occipital gyrus and fusiform gyrus. The outcome of Step 1 was therefore a time × time matrix that represented the contribution of occipito-temporal sources to the decoding model.

Step 2: To consider both the decoding strength and localization strength of each source, we computed source representation as the dot product between the time × time decoding performance (after statistical test, see *Top-Down Cross-decoding Step 2*) and the time × time source localization (see *Step 1*).

Step 3: We further down-sampled time × time representation for each source just as we did in *Top-Down Cross-decoding Step 3* to capture the source representation of the early-late bottom-up × early-late top-down decoding.

Step 4: We applied 2 (left vs. right-located prediction) * 2 (left vs. right hemisphere) ANOVA on the late top-down representation (decoded by early bottom-up) on occipital sources to test the interaction effect (i.e., the contra-lateral effect).

We repeated these four steps for each participant, creating per participant the down-sampled 2-by-2 matrix to show this source representation of early top-down decoded by late bottom-up and late top-down decoded by early bottom-up (see Supplemental Figure S2 for each individual result; and Supplemental Table S4 for each individual result). Figure 2 shows the group average source representation, separately for left and right located predictions.

#### Representation Overlap

To determine whether trial-per-trial Gabor predictions at Stage 2 comprise the same SF contents as when the Gabor stimuli are actually processed at Stage 3, we quantified the overlap of Gabor LSF vs. HSF representations between Stages 2 and 3. We did so with information theoretic redundancy quantified with co-information [41, 42] —i.e. Co-I(<LSF vs. HSF, Stage 2 MEG; Stage 3 MEG>) on trials with valid predictions and separately for left and right localized stimuli. This quantifies the information about LSF vs HSF which is shared between or common to the Stage 2 and Stage 3 MEG responses.

Step 1: To simplify calculations, we chose one source at Stage 2: one for the contra-lateral occipital cortex and one for the right fusiform gyrus. In each region, we chose that source that maximizes MI(<LSF vs. HSF prediction; Stage 2 MEG>) computed every 2ms over the [0 - 500ms] interval post auditory cue.

Step 2: To trace the per-trial representational overlap between Gabor LSF vs. HSF predictions at Stage 2 and actual LSF vs. HSF Gabor representations at Stage 3, we used 6 ROIs—i.e. (1) contra-lateral occipital cortex, (2) right fusiform gyrus, (3) temporal lobe (inferior temporal gyrus, middle temporal gyrus, superior temporal gyrus), (4) parietal lobe (inferior parietal lobe, superior parietal lobe, supramarginal gyrus and angular gyrus), (5) frontal lobe (orbitofrontal cortex, inferior frontal gyrus, middle frontal gyrus, medial frontal gyrus and superior frontal gyrus), (6) pre-motor cortex (precentral gyrus and postcentral gyrus).

Step 3: We then computed the single-trial information theoretic redundancy— i.e. the Co-Information Co-I(<Stage 2 MEG; Stage 3 MEG; LSF vs. HSF>)

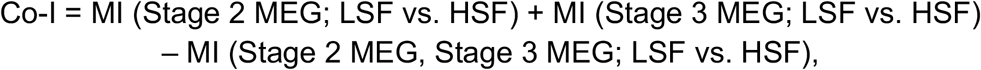

where Stage 2 MEG refers to the single-trial activity of the 2 sources chosen at Step 1 above; Stage 3 MEG refers to each source of the 6 ROIs defined in Step 2 above. We computed Co-I on the full combinatorics of Stage 2 MEG and Stage 3 MEG, every 2ms between 0 and 500ms post Gabor onset, using valid informative cueing trials when the single-trial LSF vs. HSF label is consistent between Stages 2 and 3. This computation produced a 4D matrix (Stage 2 sources × Stage 3 sources × Stage 2 time × Stage 3 time).

Step 4: When the Gabor appears at Stage 3, we quantified how its representation in each one of the 6 ROIs overlaps with its prediction at Stage 2. In each ROI, we selected the top 25% of the maximum Co-I values (taken for each source, at peak Co-I time at Stage 3) and averaged these values per ROI. This produced a 2×6 matrix for Co-I peak amplitudes and peak times (i.e. 2 sources selected at Stage 2 × the top 25% Co-I in each ROI at Stage 3).

We repeated the above Steps 1-4 for each participant. Round and diamond scatters above plots A and B in Figure 3 show the individual Co-I peak amplitude and peak time at Stage 3 in contra-lateral occipital cortex, separately for right and left-located trials. Supplemental Figure S3 Y-axis shows the individual-level and group-level Co-I strength for all 6 ROIs at Stage 3.

#### Cue modulates Source Activity and RT

To detect how valid predictions modulate MEG activity and facilitate behavior, we compared the effect of Stage 2 prediction (i.e., predicted vs. non-predicted) jointly on Stage 3 brain activity and RT.

Step 1: We computed single-trial information theoretic redundancy—i.e. positive Co-I(<predicted vs. non-predicted; Stage 3 MEG; RT>)—as follows:

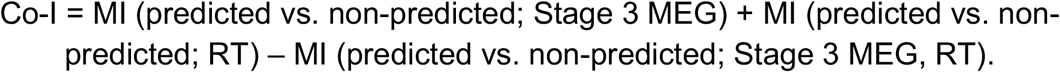

for all sources of the 6 ROIs at Stage 3 (see <u>Representation Overlap</u> Step 2), every 2ms between 0 and 500ms post Gabor onset. Co-I computations produced a source-by-time redundancy matrix. We established statistical significance by recomputing the matrix with shuffled cueing labels (repeated 1,000 times), using as statistical threshold the 95^th^ percentile of 1,000 maxima (each taken across the redundancy matrix of each shuffled repetition, FWER, *p*<0.05, one-tailed).

Step 2: In each ROI, we quantified how the valid informative cueing modulated brain activity and RT. We averaged the top 25% of maximum Co-I values in each ROI (taken for each source, across time, after statistical test) at their peak time. This produced a vector of 6-ROI redundancy peak amplitudes and peak times.

We applied Step 1 and 2 to each participant, to derive peak values of redundancy strength and peak time for each of 6 ROIs. The square scatters above plots C and D in Figure 3 show the Co-I peak time and amplitude in pre-motor cortex, separately for right and left located trials. Supplemental Figure S3 X-axis shows the individual-level and group-level redundancy strength in all 6 ROIs at Stage 3.

#### Prediction Facilitation on SF Representation

To understand how Stage 2 predictions facilitate Stage 3 visual categorizations, we compared representation of Gabor LSF vs. HSF between predicted and non-predicted trials at Stage 3. Specifically, we computed MI between MEG source and LSF vs. HSF with predicted (valid informative cued) vs. non-predicted (neutral cued) trials in contra-lateral occipital cortex, to quantify the facilitation effect on early visual processing (Step 1) and also in premotor cortex, to quantify the facilitation effect on later decision making (Step 2), separately for left and right located trials:

Step 1: On contra-lateral occipital sources with top 25% representation overlap (see <u>Representation Overlap</u> Step 5), we computed source-by-time MI(LSF vs. HSF; Stage 3 MEG), every 2ms between 0 and 500ms post Gabor onset, separately for predicted and non-predicted trials, matching trial numbers with random selection for this computation and averaging 5 random trial selections. We established the statistical significance of comparisons between predicted MI and non-predicted MI by recomputing the source-by-time MI with shuffled predicted and unpredicted trials (repeated 1,000 times), calculating the difference between recomputed predicted and non-predicted MI per time point per source, using as statistical threshold the 95^th^ percentile of 1,000 maxima, each taken across the source-by-time difference of each shuffled repetition (FWER, *p*<0.05, two-tailed). Separately for predicted and non-predicted trials, across those sources, we averaged MI with significant difference within the 50ms time window following their peak representational overlap (see <u>Representation Overlap</u> Step 5).

Step 2: We computed MI(LSF vs. HSF; Stage 3 MEG) as in Step 1, this time using pre-motor sources with top 25% Co-I (predicted vs. non-predicted; Stage 3 MEG; RT) (see <u>Cue modulates Source Activity and RT</u> Step 2), applying statistical testing as in Step 1, averaging source MI with significant difference within the 50ms time window after their peak time of Co-I (predicted vs. non-predicted; Stage 3 MEG; RT) (see <u>Cue Modulates Source Activity and RT</u> Step 2).

We repeated above Step 1-2 for each participant. In Figure 3, curves A-D show the averaged MI time course across sources from a typical participant. Supplemental Figure S4 scatters the mean SF representation across the 50ms time window for each participant, separately for right and left-located trials.

## Acknowledgements

P.G.S. received support from the Wellcome Trust (Senior Investigator Award, UK; 107802) and the Multidisciplinary University Research Initiative/Engineering and Physical Sciences Research Council (USA, UK; 172046-01). R.A.A.I. was supported by the Wellcome Trust [214120/Z/18/Z]. The funders had no role in study design, data collection and analysis, decision to publish or preparation of the manuscript.

## Author contributions

Y.Y., J.Z., P.G.S designed the research; Y.Y., performed the experiment; Y.Y. and J.Z. analyzed the data; R.A.A.I and P.G.S gave suggestions on data analysis; and Y.Y., J.Z., R.A.A.I, and P.G.S wrote the paper.

