## Supplemental Materials for "Top-down predictions of visual features dynamically reverse their bottom-up processing in the occipito-ventral pathway to facilitate stimulus disambiguation and behavior"

**Supplemental Table S1.** Group-level effect of cueing on mean Gabor categorization RTs (for left vs. right located, LSF vs. HSF), tested with paired samples t-test.

| Gabor type | RT (predicted, ms) | RT (non-predicted, ms) | Improvement on RT (ms) | *t* value | *p* value |
|---|---|---|---|---|---|
| Left LSF | 530.9 | 456.8 | 74.1 | $t_{(10)} = 3.60$ | $p = 0.005$ |
| Left HSF | 555.4 | 447.7 | 107.7 | $t_{(10)} = 5.87$ | $p = 0.0002$ |
| Right LSF | 556.3 | 483.6 | 72.6 | $t_{(10)} = 3.37$ | $p = 0.007$ |
| Right HSF | 525.9 | 429.5 | 96.4 | $t_{(10)} = 4.82$ | $p = 0.0007$ |

**Supplemental Table S2.** Individual-level effect of predicted vs. non-predicted on categorization RTs of Gabor patches (2 (left vs. right-located) x 2 (LSF vs. HSF) conditions, tested with ANOVA and independent samples *t*-test (before Bonferroni correction).

| Participant ID | ANOVA *F*-value: Effect of valid informative vs. neutral cueing | Left LSF mean RT reduction | Right LSF mean RT reduction | Left HSF mean RT reduction | Right HSF mean RT reduction |
|---|---|---|---|---|---|
| 1 | $F(1, 2174) = 2765.5$, $p<0.001$ | 71.7ms, $p<0.001$ | 87.8ms, $p<0.001$ | 127.4ms, $p<0.001$ | 121.9ms, $p<0.001$ |
| 2 | $F(1, 1968) = 90.1$, $p<0.001$ | 42.9ms, $p<0.001$ | 30.0ms, $p=0.002$ | 57.8ms, $p<0.001$ | 45.0ms, $p<0.001$ |
| 3 | $F(1, 1888) = 133.5$, $p<0.001$ | 30.2ms, $p=0.008$ | 31.8ms, $p<0.001$ | 96.9ms, $p<0.001$ | 103.0ms, $p<0.001$ |
| 4 | $F(1, 2147) = 75.1$, $p<0.001$ | **-2.6ms, *p*=0.71** | **1.2ms, *p*=0.87** | 65.2ms, $p<0.001$ | 59.1ms, $p<0.001$ |
| 5 | $F(1, 1889) = 42.3$, $p<0.001$ | **16.8ms, *p*=0.12** | **19.7ms, *p*=0.04** | 44.6ms, $p<0.001$ | 38.5ms, $p<0.001$ |
| 6 | $F(1, 1751) = 237.6$, $p<0.001$ | 139.3ms, $p<0.001$ | 115.2ms, $p<0.001$ | 102.2ms, $p<0.001$ | 136.9ms, $p<0.001$ |
| 7 | $F(1, 1685) = 241.7$, $p<0.001$ | 81.5ms, $p<0.001$ | 111.5ms, $p<0.001$ | 127.6ms, $p<0.001$ | 93.8ms, $p<0.001$ |
| 8 | $F(1, 1704) = 1887.1$, $p<0.001$ | 149.5ms, $p<0.001$ | 173.1ms, $p<0.001$ | 234.2ms, $p<0.001$ | 238.5ms, $p<0.001$ |
| 9 | $F(1, 1806) = 28.0$, $p<0.001$ | **26.2ms, *p*=0.046** | **18.2ms, *p*=0.15** | 58.6ms, $p<0.001$ | 35.8ms, $p=0.007$ |
| 10 | $F(1, 1510) = 31.0$, $p<0.001$ | **15.1ms, *p*=0.34** | **55.9ms, *p*=0.016** | 108.2ms, $p<0.001$ | **27.5ms, *p*=0.10** |
| 11 | $F(1, 1786) = 2424.5$, $p<0.001$ | 176.3ms, $p<0.001$ | 162.9ms, $p<0.001$ | 139.9ms, $p<0.001$ | 135.1ms, $p<0.001$ |

**Bold** indicates **non-significant** result ($p>0.05$, Bonferroni-corrected).

**Supplemental Table S3.** Individual-level effect of predicted vs. non-predicted on categorization accuracy Gabor patches (2 (left vs. right-located) x 2 (LSF vs. HSF) conditions, tested with ANOVA and chi-squared tests (before Bonferroni correction).

| Participant ID | ANOVA *F*-value: Effect of valid informative vs. neutral cueing | Left LSF mean ACC increase | Right LSF mean RT increase | Left HSF mean RT increase | Right HSF mean RT increase |
|---|---|---|---|---|---|
| 1 | ***F*(1, 2248) = 7.24, *p*=0.007** | 0% | -0.26%, *p*=0.47 | **1.7%, *p*=0.011** | 1.1%, *p*=0.04 |
| 2 | *F*(1, 2098) = 0.06, *p*=0.80 | 0.15%, *p*=0.85 | 1.5%, *p*=0.31 | -2.3%, *p*=0.04 | 0.08%, *p*=0.94 |
| 3 | ***F*(1, 1998) = 10.2, *p*<0.001** | 1.7%, *p*=0.19 | **3.7%, *p*=0.006** | 2.0%, *p*=0.23 | 1.6%, *p*=0.20 |
| 4 | ***F*(1, 2348) = 11.8, *p*<0.001** | -1.5%, *p*=0.37 | 2.6%, *p*=0.27 | 6.8%, *p*=0.014 | **7.2%, *p*<0.001** |
| 5 | ***F*(1, 2048) = 7.33, *p*=0.007** | -0.23%, *p*=0.91 | **7.7%, *p*=0.006** | 1.0%, *p*=0.59 | **2.7%, *p*=0.011** |
| 6 | ***F*(1, 1948) = 21.3, *p*<0.001** | 4.0%, *p*=0.11 | 6.2%, *p*=0.013 | **7.4%, *p*<0.001** | 3.7%, *p*=0.08 |
| 7 | ***F*(1, 1848) = 14.2, *p*<0.001** | -1.1%, *p*=0.60 | 9.9%, *p*=0.003 | **7.5%, *p*<0.001** | 1.6%, *p*=0.08 |
| 8 | ***F*(1, 1898) = 13.0, *p*<0.001** | 1.6%, *p*=0.31 | 2.2%, *p*=0.32 | **10%, *p*<0.001** | 2.1%, *p*=0.30 |
| 9 | *F*(1, 1948) = 1.09 *p*=0.30 | -1.5%, *p*=0.13 | 0.6%, *p*=0.71 | 1.7%, *p*<0.001 | 2.5%, *p*=0.007 |
| 10 | *F*(1, 1798) = 0.96, *p*=0.32 | -1.2%, *p*=0.34 | 0.83%, *p*=0.016 | 4.1%, *p*=0.44 | -0.02%, *p*=0.08 |
| 11 | ***F*(1, 1898) = 36.6, *p*<0.001** | **3.9%, *p*<0.001** | **3.4%, =0.005** | **3.3%, *p*=0.008** | **2.8%, *p*=0.002** |

**Bold** indicates **significant** (*p*<0.05 after Bonferroni-correction) result.

**Supplemental Table S4.** Individual-level contra-lateral effect of source representation of LSF vs. HSF prediction, (left vs. right-predicted location) x 2 (left vs. right occipital sources), tested with ANOVA.

| Participant ID | ANOVA *F*-value: Effect of predicted location | ANOVA *F*-value: Effect of hemisphere | ANOVA *F*-value: Effect of predicted location*hemisphere |
|---|---|---|---|
| 1 | $F(1, 1308) = 4.31$, $p=0.29$ | $F(1, 1308) = 1.00$, $p=0.50$ | **$F(1, 1308) = 565.7$, $p<0.001$** |
| 2 | $F(1, 1308) = 0.84$, $p=0.53$ | $F(1, 1308) = 0.48$, $p=0.62$ | **$F(1, 1308) = 181.7$, $p<0.001$** |
| 3 | $F(1, 1308) = 1.52$, $p=0.43$ | $F(1, 1308) = 0.63$, $p=0.57$ | **$F(1, 1308) = 71.4$, $p<0.001$** |
| 4 | NaN | NaN | NaN |
| 5 | $F(1, 1308) = 2.16$, $p=0.38$ | $F(1, 1308) = 1.08$, $p=0.49$ | **$F(1, 1308) = 77.9$, $p<0.001$** |
| 6 | $F(1, 1308) = 0.25$, $p=0.71$ | $F(1, 1308) = 0.72$, $p=0.55$ | **$F(1, 1308) = 166.5$, $p<0.001$** |
| 7 | NaN | NaN | NaN |
| 8 | $F(1, 1308) = 1.35$, $p=0.45$ | $F(1, 1308) = 1.00$, $p=0.50$ | **$F(1, 1308) = 169.9$, $p<0.001$** |
| 9 | $F(1, 1308) = 12902$, $p=0.005$ | $F(1, 1308) = 172.8$, $p=0.05$ | $F(1, 1308) = 0.05$, $p=0.82$ |
| 10 | $F(1, 1308) = 2.51$, $p=0.36$ | $F(1, 1308) = 1.33$, $p=0.46$ | **$F(1, 1308) = 60.7$, $p<0.001$** |
| 11 | $F(1, 1308) = 17.6$, $p=0.15$ | $F(1, 1308) = 2.5$, $p=0.36$ | **$F(1, 1308) = 36.6$, $p<0.001$** |

**Bold** indicates **significant interaction effect.**

**Supplemental Table S5.** Individual-level contrast and accuracy of LSF vs. HSF Gabor patches in contrast threshold testing.

| Participant ID | Left LSF Contrast (ACC) | Right LSF Contrast (ACC) |
|---|---|---|
| 1 | 0.2277 (91.5%) | 0.1752 (91.9%) |
| 2 | 0.2152 (91.1%) | 0.1752 (91.9%) |
| 3 | 0.2021 (92.3%) | 0.1720 (91.0%) |
| 4 | 0.2215 (91.3%) | 0.2102 (92.5%) |
| 5 | 0.1633 (93.0%) | 0.1815 (91.4%) |
| 6 | 0.2019 (92.9%) | 0.1852 (91.7%) |
| 7 | 0.2219 (91.5%) | 0.1952 (91.5%) |
| 8 | 0.2367 (83.8%) | 0.1967 (94.6%) |
| 9 | 0.2300 (92.6%) | 0.2152 (92.0%) |
| 10 | 0.2356 (90.5%) | 0.2252 (92.7%) |
| 11 | 0.2152 (91.1%) | 0.1668 (92.7%) |

**Supplemental Table S6.** Stimulus repetition in one cueing-categorization block

| | Location cue | SF cue | Visual stimuli (random from 3 orientations) |
|---|---|---|---|
| Repetitions/type | 27 left cues | 9 LSF cues | 8 left-LSF + 1 left-HSF |
| | | 9 HSF cues | 8 left-HSF + 1 left-LSF |
| | | 9 neutral cues | 9 left-random LSF/HSF |
| | 27 right cues | 9 LSF cues | 8 right-LSF + 1 right-HSF |
| | | 9 HSF cues | 8 right-HSF + 1 right-LSF |
| | | 9 neutral cues | 9 right-random LSF/HSF |
| Sum | 54 | | |

**Supplemental Table S7.** Trials remaing following pre-processing and LCMV analysis

| Stage | Mean trial number/participant | Range |
|---|---|---|
| Stage 1 | 1609 | 1492-1759 |
| Stage 2 | 1604 | 1505-1751 |
| Stage 3 | 1620 | 1507-1743 |
| Localizer | 592 | 336-699 |

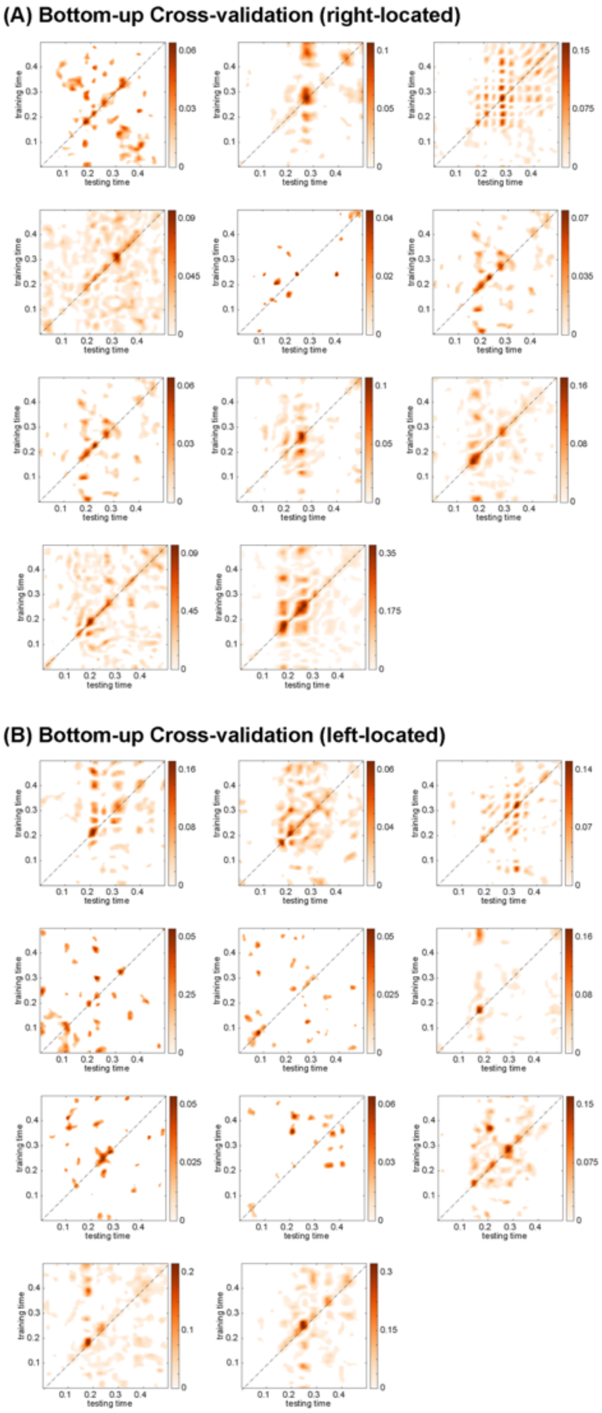


**Supplemental Figure S1 Bottom-up cross-validation performance at individual-level for (A) right-located and (B) left-located trials.** We computed and tested Gabor LSF vs. HSF classifiers on the localizer dataset by applying a 5-fold cross-validation with 10 repetitions and quantified the cross-validation performance by computing MI between decision values and true “LSF” vs. “HSF” labels. Each color-indexed matrix represents significant cross-validation performance (MI/bit, FWER, $p < 0.05$, after TFCE, one-tailed) in each participant, demonstrating that bottom-up SF classifiers decoded top-down predicted LSF vs. HSF contents.

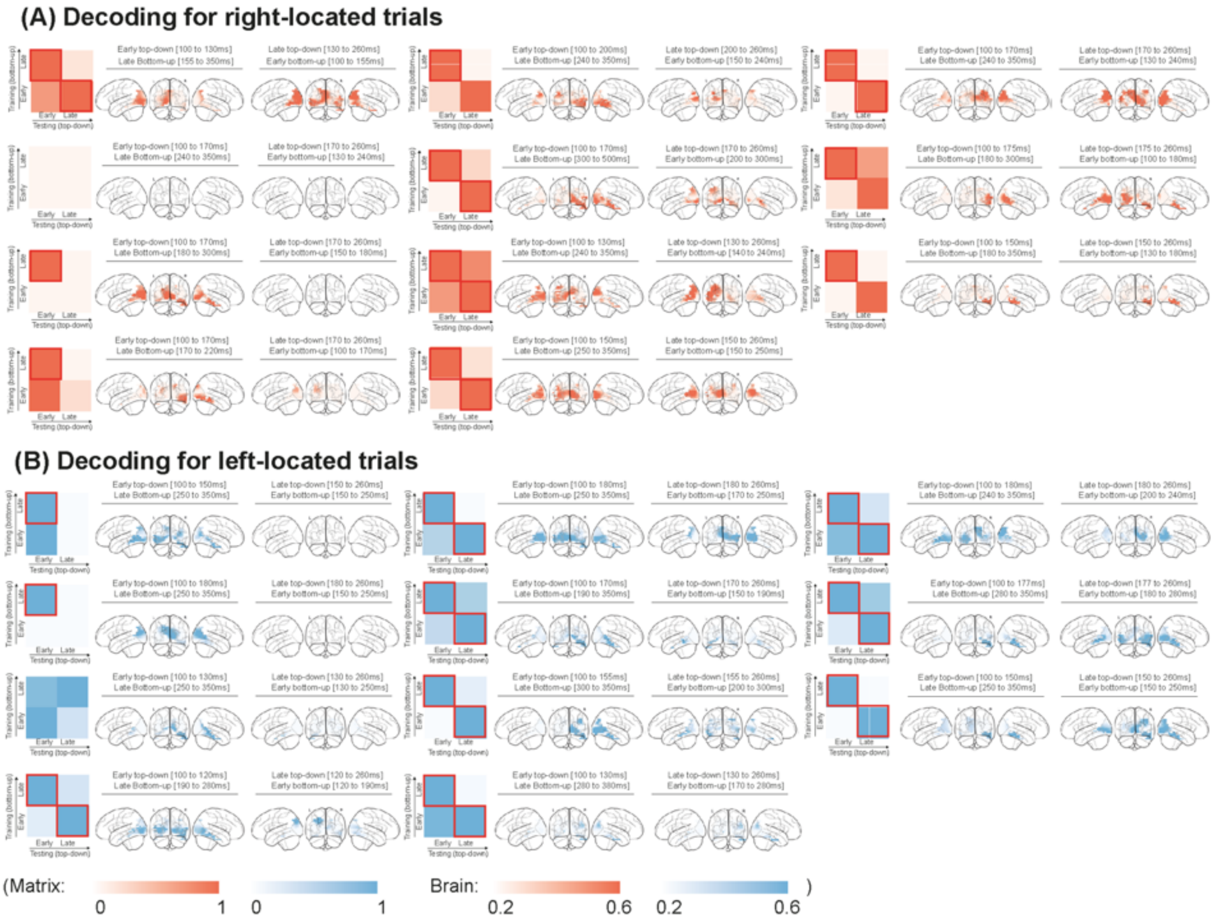


**Supplemental Figure S2 Decoding performance and source representation at Individual-level for (A) right-located and (B) left-located trials.** Matrices. For right (color-coded in orange) and left (color-coded in blue) located Gabor trials, each matrix illustrates the decoding performance obtained in each participant. We used the bottom-up SF classifier (learned from MEG localizer) to decode the LSF vs. HSF contents of Stage 2 top-down predictions (MEG sensor data). We quantified decoding performance as MI(<decoder value; ground truth LSF vs. HSF>). We down-sampled the significant decoding performance (FWER, $p<0.05$, one-tailed) into a 2-by-2 matrix that averaged each MI matrix along the Y-axis that represents the bottom-up time window (i.e., early – late localizer; training dataset) and X-axis that represents the top-down time window (i.e., early – late prediction; testing dataset), with the specific time windows indicated above the brain plots. We normalized within each classifier (i.e., normalized across each row) to highlight that the early bottom-up classifier better decodes the late top-down responses, and the late bottom-up classifier better decodes the early top-down responses. Brain plots show the Stage 2 source representation of significant Gabor LSF vs. HSF decoding. We first calculated MI(<decoder value; Stage 2 MEG) along the occipito-ventral pathway (i.e. lingual gyrus, cuneus, inferior occipital gyrus, middle occipital gyrus, superior occipital gyrus and fusiform gyrus), and then computed source representation as the dot product between the time × time decoding performance and the time × time MI(decoder value; Stage 2 MEG). Similarly, we down-sampled the source representation with the early-late top-down × early-late bottom-up time window. We showed the source representation of early top-down decoded by late bottom-up (left-side brain plots) and late top-down decoded by early bottom-up (right-side brain plots), normalized across classifiers, revealing the top-down flow from right fusiform gyrus down to the primary occipital cortex contra-lateral to the predicted location of the upcoming Gabor.

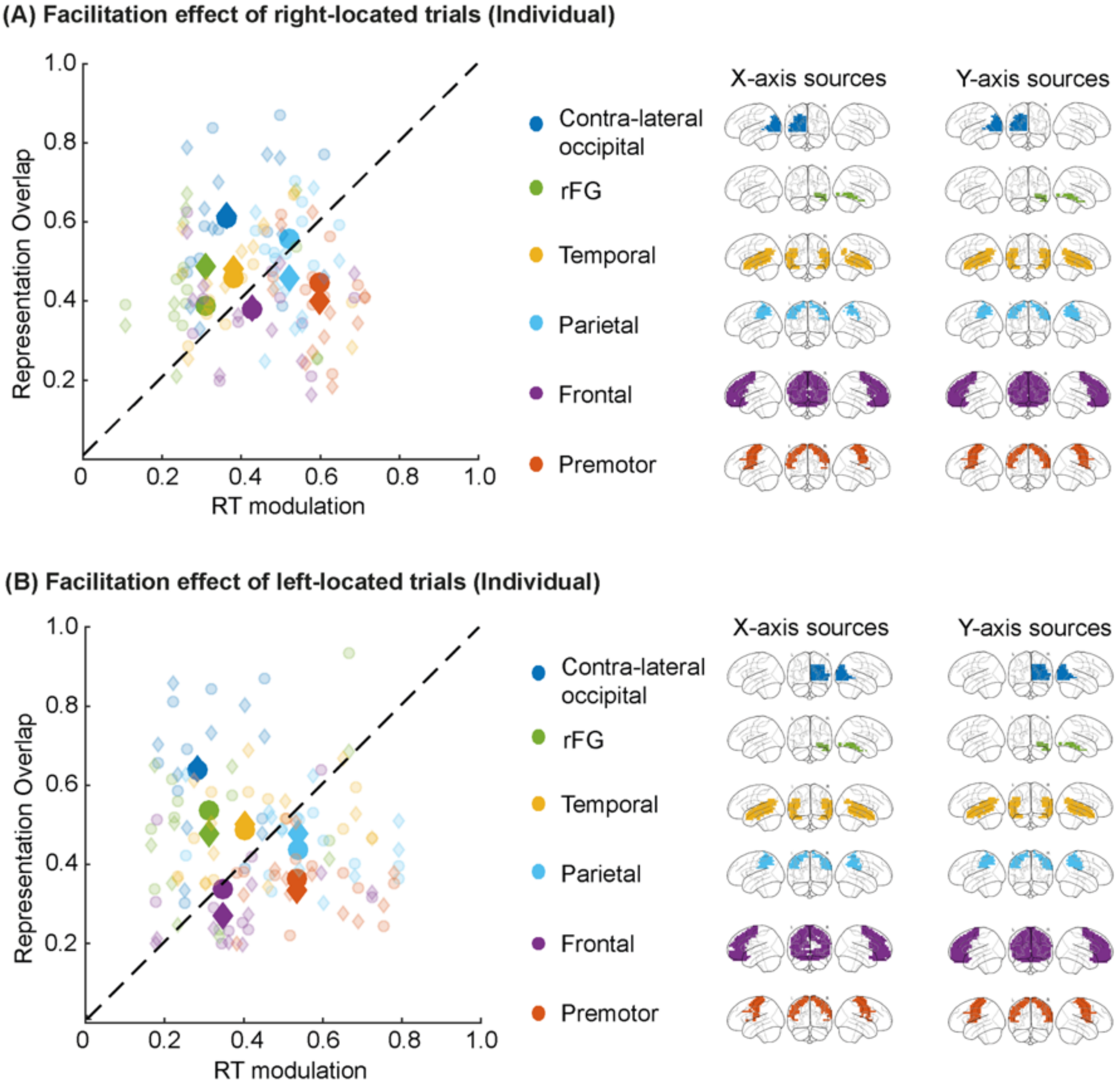


**Supplemental Figure S3 SF redundancy – RT redundancy distribution across the whole brain for right-located (A) and left-located (B) trials.** Left panel shows 3.1 Co-I(LSF vs. HSF; Stage 2 MEG; Stage 3 MEG) on the X-axis, 3.2 Co-I(predicted vs. non-predicted; RT; Stage 3 MEG) on the Y-axis. Color-indexed scatters show SF redundancy (X-axis, circles, redundancy with Stage 2 contra-lateral occipital source, diamonds, redundancy with Stage 2 right fusiform gyrus source), RT redundancy (Y-axis) in contra-lateral occipital cortex (blue), right fusiform gyrus (green), temporal lobe (yellow, including inferior temporal gyrus, middle temporal gyrus and superior temporal gyrus), parietal lobe (light blue, including inferior parietal lobe, superior parietal lobe, supramarginal gyrus and angular gyrus), frontal lobe (purple, including orbitofrontal cortex, inferior frontal gyrus, middle frontal gyrus, medial frontal gyrus and superior frontal gyrus), and premotor cortex (orange, including precentral gyrus and postcentral gyrus). Each t scatter represents the mean of top 25% of maximum Co-I (LSF vs. HSF; Stage 2 MEG; Stage 3 MEG) value (taken for each Stage 3 source, across Stage 2 times x Stage 3 times), the mean value of top 25% of maximum Co-I (predicted vs. non-predicted; RT; Stage 3 MEG) value (taken for each Stage 3 source, across Stage 3 times). Each opaque scatter represents the group-level mean result. Right panel shows the location of color-coded ROI sources that have significant Co-I in the top 25% of maximum Co-I for at least one of the participants.

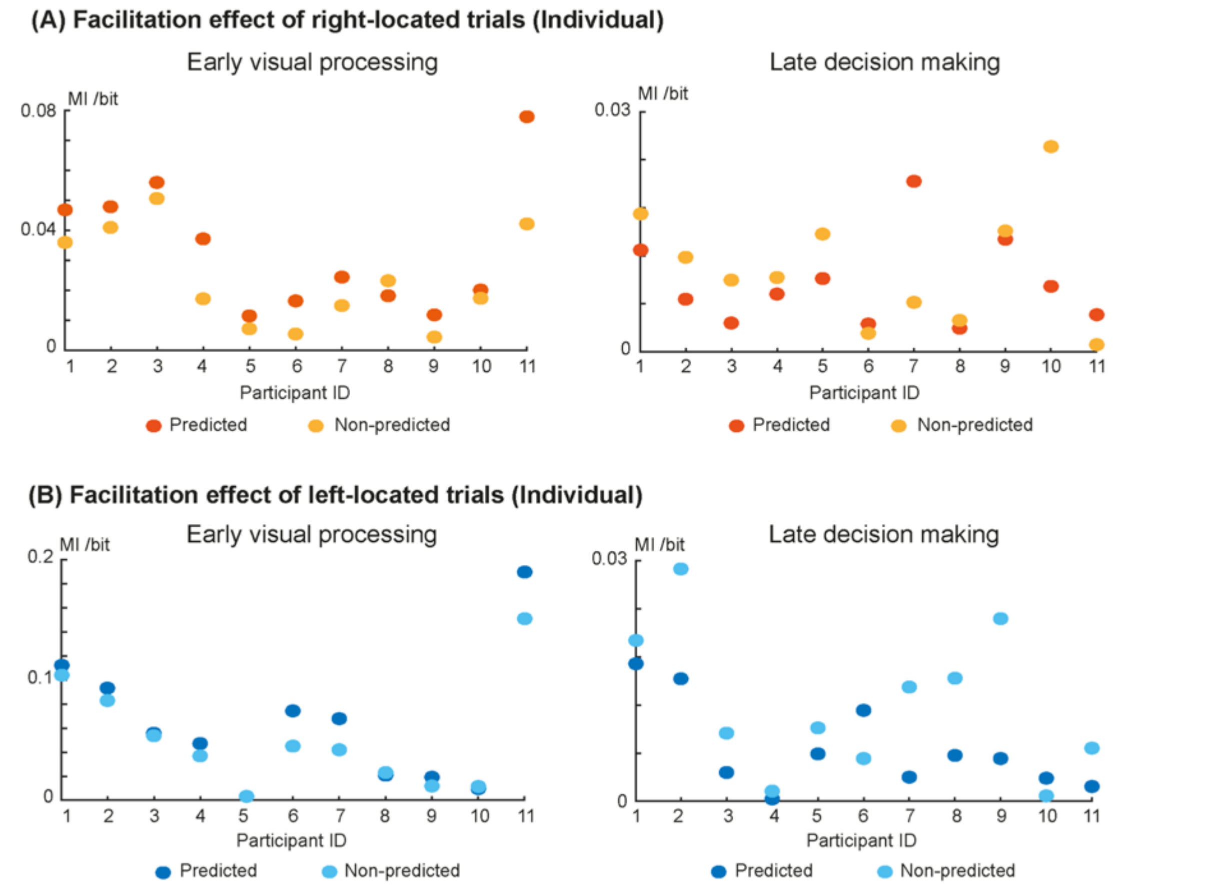


**Supplemental Figure S4 Facilitation of Gabor LSF vs. HSF source representations at individual level for left-located (A) trials and right-located (B) trials.** In (A) and (B), the blue (vs. orange) scatters show the mean representation of Gabor LSF vs. HSF, computed as MI(LSF vs. HSF; Stage 3 MEG) (Y-axis) in left- (vs. right-) located trials. Left panel. Across contra-lateral occipital sources at the top 25% of the Co-I(LSF vs. HSF; Stage 2 MEG; Stage 3 MEG) distribution, we averaged the SF representation from 0 to 50 ms following the Co-I peak time. Darker (vs. lighter) hues represent predicted (vs. non-predicted) trials. X-axis numbers represent each participant. 8/11 (vs. 10/11) participants had significantly improved LSF vs. HSF discrimination on valid informative cueing in left-located (vs. right-located) trials (FWER, $p<0.05$, two-tailed). Right panel Across pre-motor cortex sources at the top 25% of the Co-I(predicted vs. non-predicted; Stage 3 MEG; RT) distribution, we averaged the SF representation from 0 to 50 ms following the Co-I peak time. Darker (vs. lighter) hues represent predicted (vs. non-predicted) trials. X-axis numbers represent each participant. 9/11 (vs. 8/11) participants had significantly decreased representation of SF discrimination on valid informative cueing in left- (vs. right-) located trials.
